# The hyaluronan receptor CD44 drives COVID-19 severity through its regulation of neutrophil migration

**DOI:** 10.1101/2025.10.13.682000

**Authors:** Duncan J. Hart, Md. Jashim Uddin, Rebecca J. Dodd, Savannah G. Brovero, Claire Fleming, G. Brett Moreau, Nick R. Natale, Barbara J. Mann, Tara E. Sutherland, Judith E. Allen, Anthony J. Day, William A. Petri

## Abstract

The novel respiratory disease COVID-19 caused by the coronavirus SARS-CoV-2 continues to be a public health emergency worldwide, and there is a need for more effective therapy for patients. The relationship between the extracellular matrix and the host immune response to infection is severely understudied. Deposition of the polysaccharide hyaluronan (HA) into the lungs is associated with more severe COVID-19 disease outcomes. HA is a major component of the extracellular matrix in connective tissues and is abundant in many parts of the body, including cartilage, skin, brain, and vitreous body. CD44 is the primary receptor for HA and is found on almost all immune cells in the lung. Known functions of CD44 include mediation of immune cell migration, activation, and differentiation. We hypothesized that increased HA deposition during COVID-19 increases CD44-mediated immune cell infiltration into lungs and results in more severe pathology. Here, we report that in mice infected with a mouse-adapted strain of SARS-CoV-2, treatment with a combination of two anti-CD44 monoclonal antibodies confers a significant survival benefit and reduces weight loss and clinical score of the mice on Day 4 post infection. We show that anti-CD44 treatment decreases many key cytokines and chemokines in the bronchoalveolar lavage fluid on Day 4. With flow cytometry, we show that anti-CD44 reduces the numbers of neutrophils in infected lungs. We also show through immunofluorescence that treatment with anti-CD44 antibodies reduces colocalization of HA and CD45 in lung sections, indicating that HA’s interaction with immune cells contributes to pathology. Our findings demonstrate that disruption of HA-receptor interactions is a way to prevent inflammatory pathology in pulmonary infection.

## Introduction

Infection with SARS-CoV-2 has become endemic, and there remains a need for more effective therapy especially for patients with severe COVID-19. Pulmonary production of the polysaccharide hyaluronan (HA) is associated with more severe COVID-19 disease outcomes [1–6]. HA is a glycosaminoglycan that is a key component of the extracellular matrix through its interaction with proteins and proteoglycans.[7] HA is produced throughout the body by a variety of cell types and is involved in many immune pathways [8–10]. HA forms a pathogenic matrix that retains water and promotes the adhesion of leukocytes [11–13] in patients with asthma [14] and in mouse models of influenza [15] and *Nippostrongylus brasiliensis* infection [16]. These viscous ‘gels’ have also been found in severe COVID-19 patients [5] with respiratory distress syndrome (ARDS)-like symptoms [13,17] and can be simulated by hypoxia [18]. In addition, HA is implicated in inflammation in inflammatory bowel disease [9,19,20]. The properties of HA are determined in part by the number of disaccharide repeats present as well as by HA-binding proteins [7]. For example, low molecular weight (LMW) HA fragments from severe COVID-19 patients have been found in vitro to activate alveolar macrophages and induce the release of inflammatory mediators, negatively impacting epithelial barrier function [2]. While the molecular mechanisms underlying HA’s role in inflammation are not fully understood (and are somewhat controversial), there is strong evidence that the interaction of HA with its primary receptor CD44 plays a key role [7].

CD44 is present on many types of immune cells, including those that reside in the lungs [10,21,22]. Interactions between CD44 on immune cells and HA within the extracellular matrix and on endothelium and epithelium mediate cell adhesion and migration [23,24]. In instances of lung inflammation such as in mouse models of influenza and nematode infection, TSG-6, the secreted product of TNF-stimulated gene-6 protein [25] catalyzes the covalent transfer of the heavy chains (HC) belonging to the inter-α-inhibitor (IαI) family of proteoglycans onto HA, to form ‘HC•HA’ complexes [15,16,26]. Subsequent interactions between the covalently bound HCs, and also with the octameric protein pentraxin-3, lead to the association of HA chains, and the formation of a crosslinked matrix [7,27–29]. These HC•HA complexes have been found to enhance the binding of CD44^+^ leukocytes [26,30,31] and could therefore potentially prime the lung parenchyma and airways for the infiltration of proinflammatory leukocytes in SARS-CoV-2 infection [12].

Neutrophils express high levels of CD44 [21,23,32] and are implicated in the pathogenesis of severe COVID-19 through the production of proinflammatory cytokines, reactive oxygen species, and neutrophil extracellular traps (NETs) that can elicit hypercoagulation [33–37]. In addition, cytokines and chemokines associated with neutrophil chemoattraction such as CXCL8 and IL-6 are associated with severe COVID-19 [6,38–40]. Neutrophils also bind HC•HA matrices more effectively than HA alone [26] and neutralization of CD44 has been demonstrated in animal models to mitigate neutrophil infiltration into sepsis-damaged lungs [21] and inflamed liver sinusoids [41].

In this study, we used a mouse-adapted strain of SARS-CoV-2 (MA10) to examine the role of HA-CD44 interactions in a mouse model of COVID-19. We discovered that blocking CD44 in this model suppresses lung damage. This response was at least partially mediated by a decrease in neutrophil accumulation in the lungs and a decrease in the release of inflammatory cytokines into the alveolar environment. This work indicates that disruption of HA-CD44 interactions is a way to prevent inflammatory pathology in pulmonary infection.

## Results

To test the hypothesis that hyaluronan deposition contributes to COVID-19 pathology through interactions with CD44, we intranasally inoculated mice with a mouse-adapted strain of SARS-CoV-2 that has been previously characterized [42,43]. This model replicates the age-dependent severity and much of the lung pathology of COVID-19 in humans while avoiding the extreme pathology and brain infection seen in the K18-hACE2 mouse model [44]. Mice were administered two anti-CD44 monoclonal antibodies on days 0-3 post-infection (Fig 1A). KM201 blocks the binding of CD44 to HA [45] whereas IM7 promotes shedding of CD44 from the cell surface [46,47]. We demonstrated the presence of HC•HA in lung samples of MA10 SARS-CoV-2-infected mice on day 4 post-infection, indicating the formation of covalently modified HA had occurred in our model that was not affected by combined anti-CD44 antibody treatment (Fig 1B). This combined treatment led to an overall survival benefit, less weight loss, and clinical score benefit through day 4 post-infection (Fig 1C-E). The titer of SARS-CoV-2 on day 4 post-infection was not significantly altered with treatment (Fig 1F), indicating that the protection from pathology afforded by anti-CD44 treatment is the result of changes in the host immune response rather than reduced viral load. Albumin, a known marker of lung damage [48], was measured in bronchoalveolar lavage fluid (BALF) day 4 of infection and was decreased following anti-CD44 treatment (Fig 1G). In addition, H&E staining of infected lungs showed a reduction in thickening of the alveolar wall on day 4 post-infection (Fig 1H). We concluded that the blocking of HA adhesion to CD44 in the mouse model of COVID-19 improved survival and reduced readouts associated with pulmonary immunopathology while having no impact on viral titer.

**Fig 1.**
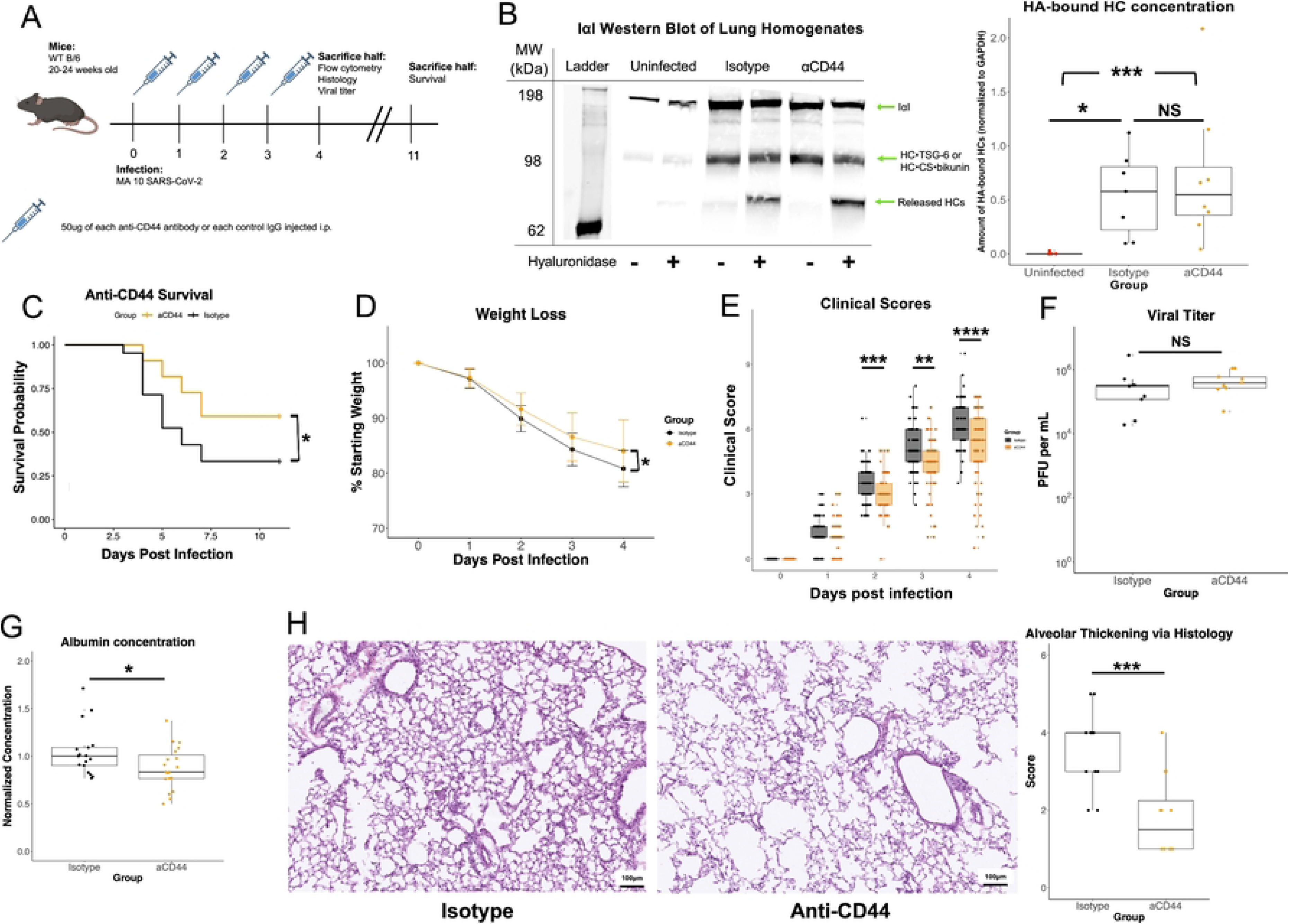
Anti-CD44 monoclonal antibody treatment reduces COVID-19 disease severity. **A.** 20–24-week-old B/6 mice were infected with 6.25 x 10^4^ PFU of MA10 SARS-CoV-2 on day 0. 50 µg of two clones of anti-CD44 mAb (aCD44; orange) or isotype controls (black) were administered intraperitoneally on days 0-3 p.i. (post infection). **B.** Representative western blotting whole lung homogenates from day 4 post infection, with/without *Streptomyces* hyaluronidase treatment, and probed with an anti-IaI antibody. HC•HA concentration (based on HCs released with hyaluronidase) were quantified via FIJI image analysis software. P values calculated via one way ANOVA followed by Tukey’s HSD test. Each data point represents one mouse. **C.** Kaplan-Meier survival analysis was done up to Day 11 post infection. **D.** Weight loss measured through day 4 p.i. Data past day 4 p.i. not shown due to survival bias. Significance calculated via linear mixed-effects model. Clinical scores of illness severity on days 0-4 p.i. Data is mean values ± SEM. **E.** Clinical scoring was measured by weight loss (0–5), posture and appearance of fur (0–2) and activity (0–3). Each data point represents one mouse. **F.** Plaque assay performed from lung homogenates taken on day 4 p.i. PFU = plaque forming units. Each data point represents one mouse. **G.** Albumin concentration measured via ELISA and normalized to control. Each data point represents one mouse. **H.** H&E stain of infected lungs on day 4 post infection. Scoring of lung damage based on alveolar thickening (0-5) was done blinded by an independent pathologist. Each data point represents one mouse. Survival, weight loss and clinical score data combined from 3 separate experiments with a total of N = 40 for each infected group, and N = 5 uninfected mice. * = p<0.05, ** = p<0.01, *** = p<0.001, **** = p<0.0001

We performed flow cytometry on whole lung samples from day 4 post-infection. Neutrophils in untreated infected lungs expressed CD44 at high levels compared to other cell types (Fig 2A). Accordingly, we observed a decrease in the number of neutrophils in whole lung samples after anti-CD44 treatment compared to isotype control (Fig 2B). IV-neutrophils were not decreased with anti-CD44 treatment compared to isotype, suggesting that anti-CD44 treatment did not kill neutrophils directly (S1 Fig). While the amounts of many other leukocytes increased with infection by varying degrees of significance, no other cell types were changed by anti-CD44 treatment (Fig 2C-D). Interestingly, B cells were unchanged with infection or treatment, indicating that B cell migration had not yet begun 4 days post infection (Fig 2D). We concluded that blockade of CD44 resulted in a decrease in infiltrating neutrophils in the lung.

**Fig 2.**
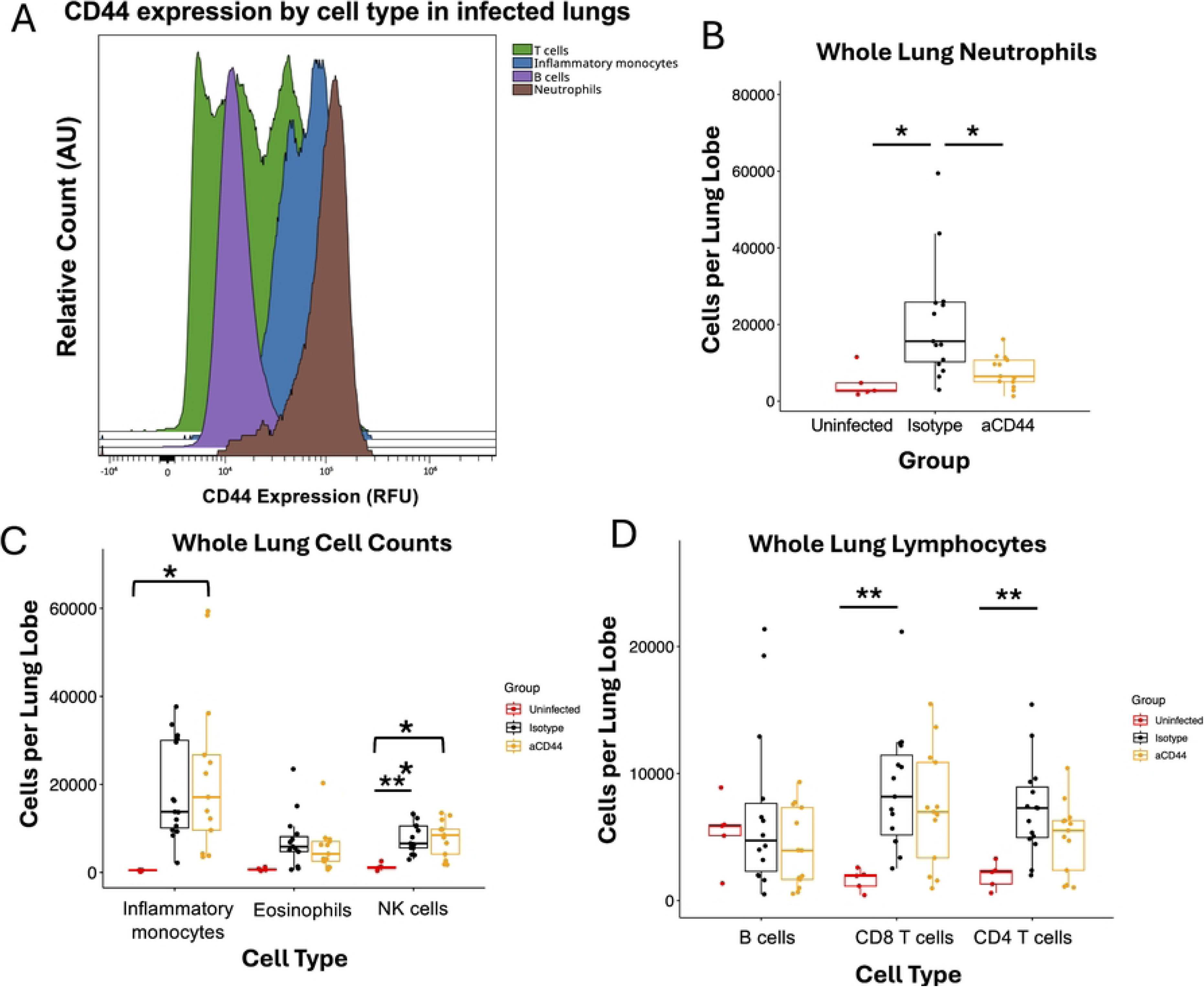
Anti-CD44 monoclonal antibody treatment reduces neutrophil infiltration into the lungs of mice during COVID-19. **A.** Histogram from flow cytometry of CD44 surface expression on white blood cells from an infected lung on day 4 post infection. AU = arbitrary units. **B.** Neutrophils (CD45^+^ CD11b^+^ CD11c^-^ Ly6C^hi^ Ly6G^+^) as count per lung lobe in whole lung homogenates taken from MA10-infected mice on day 4. **C.** Changes in white blood cell composition in whole lung homogenates taken from MA10-infected mice on day 4. Inflammatory monocytes (CD45^+^ CD11b^+^ CD11c^-^ Ly6C^hi^ Ly6G^-^), eosinophils (CD45^+^ MerTK^-^ CD64^-^ CD3^-^ NK1.1^-^ Siglec F^+^), and NK cells (CD45^+^ NK1.1^+^ SSC^lo^) are shown. **D.** Changes in whole lung lymphocyte numbers in whole lung homogenates taken from MA10-infected mice on day 4 post infection. B cells (CD45^+^ CD19^+^ SSC^lo^), CD8 T cells (CD45^+^ CD3^+^ CD8^+^ SSC^lo^), and CD4 T cells (CD45^+^ CD3^+^ CD4^+^ SSC^lo^) are shown. P values calculated via one way ANOVA followed by Tukey’s HSD test. * = p<0.05, ** = p<0.01 Relationships are not significant unless noted otherwise.

To further understand the effects that anti-CD44 treatment had on the immune response to MA-10 infection, we analyzed the levels of 32 secreted cytokines and chemokines in the BALF of infected mice on day 4 post-infection. Heatmap analysis (Fig 3A) revealed clear differences in mice treated with anti-CD44. Notably, neutrophil chemokines including CXCL1, CCL2, IL-6, and LIF were decreased upon anti-CD44 treatment (Fig 3B). IL-6 in particular is associated with severe COVID-19 in humans and has been shown to facilitate neutrophil chemoattraction via STAT3 signaling in acute inflammation [39,49]. Other chemokines directly involved in neutrophil chemotaxis such as CXCL1 and CCL2 [50,51] were also observed to have lower concentrations after anti-CD44 treatment (Fig 3B). Since these chemokines are commonly secreted by ‘activated’ epithelial cells [52,53], it is possible that the reduction in alveolar thickening seen following anti-CD44 treatment (Fig 1H) also contributes to the reduction in their concentrations in the BAL of infected mice. This indicates a possible feedback loop where neutrophils migrate to the lungs during the initial stages of SARS-CoV-2 infection, making the lung environment more pro-inflammatory, resulting in the secretion of cytokines and chemokines that then attract more neutrophils.

**Fig 3:**
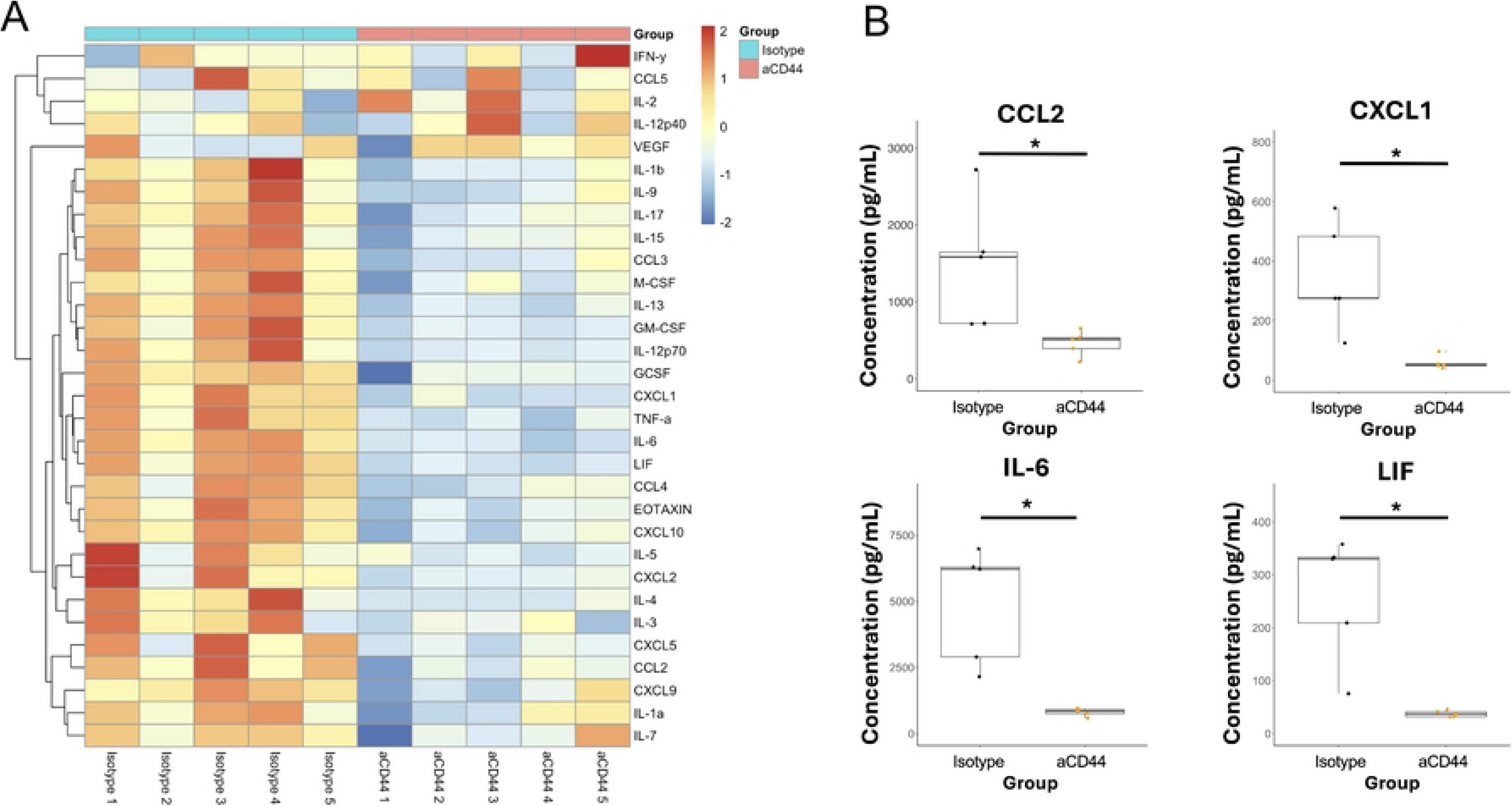
CD44 neutralization reduces the concentration of key cytokines in the BAL fluid during MA10 SARS-CoV-2 infection. **A.** Bronchoalveolar lavage fluid was collected on day 4 p.i. and protein concentrations were quantified via Luminex Multiplex analysis. Heatmap represents Z scores of log-transformed cytokine values. **B.** Key cytokines and chemokines involved in COVID-19 severity and neutrophil recruitment are shown. Data are from same analysis as in **A.** N = 5 in each group. P-values calculated with Welch’s two-sample T test * = p<0.05

To visualize the effects of anti-CD44 treatment on localization of HA and immune cells in the infected lung, we performed immunofluorescence (IF) on samples from day 4 post-infection (Fig 4A). Measured levels of CD45 and MA10 SARS-CoV-2 were not significantly different across infected groups (Fig 4B-C), aligning with our previous flow cytometry and viral titer results, respectively. Infection led to increased HA accumulation in both the airways and the alveolar spaces. Treatment with anti-CD44 significantly reduced alveolar HA, but this reduction did not reach significance in the airways. Interestingly, anti-CD44 treatment significantly reduced the colocalization of CD45^+^ cells with HA shown through two different colocalization constants (Fig 4F-G), potentially indicating a reduction in the ability of leukocytes to bind to HA matrices.

**Fig 4.**
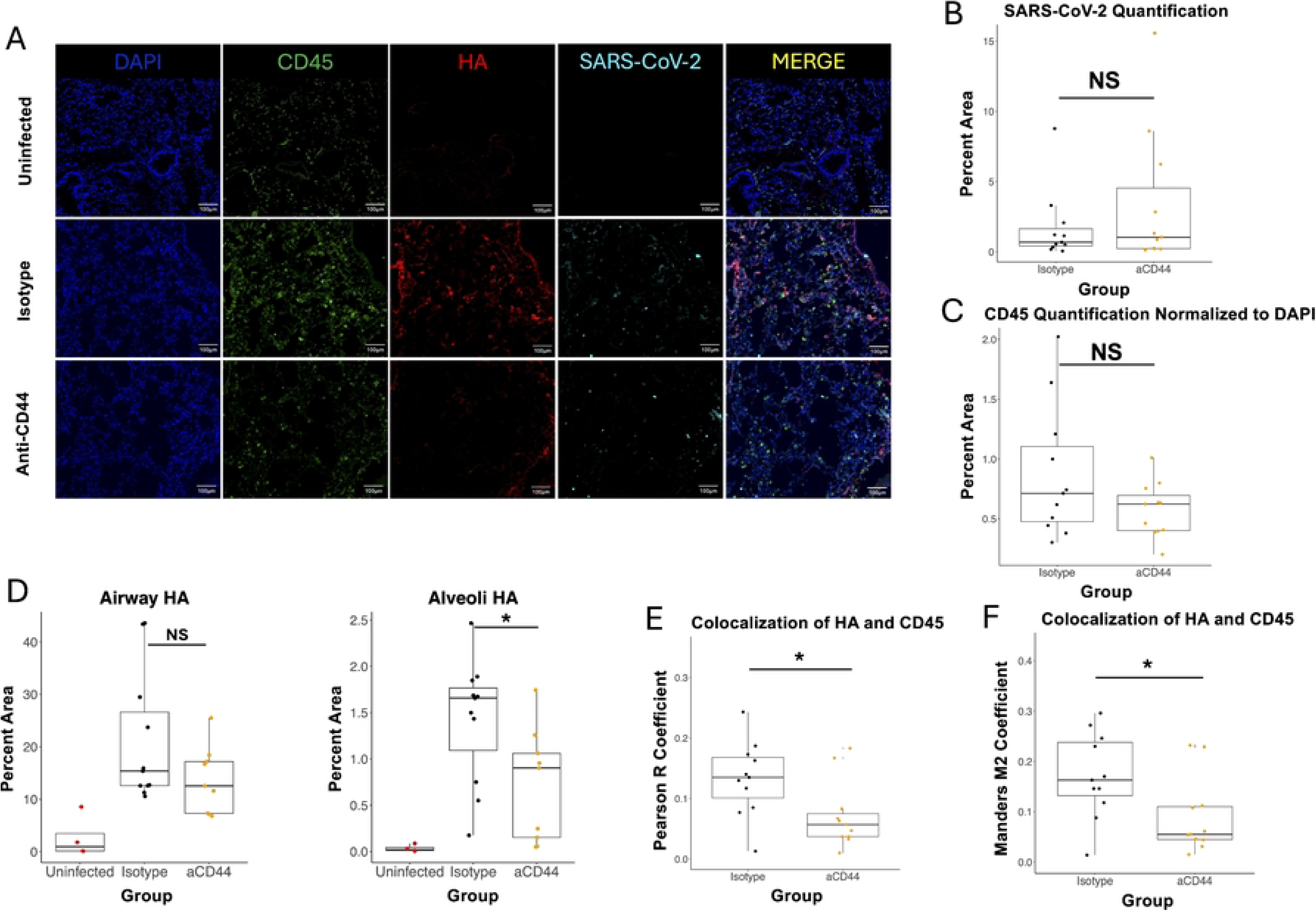
CD44 neutralization decreases colocalization between immune cells and HA. **A.** Immunofluorescence of sections from superior lung lobe section on day 4 post-infection from 20-week-old C57BL/6 mice, either uninfected or MA10-infected and treated with 50 µg of two clones of anti-CD44 mAb (aCD44) or isotype controls (Isotype) stained for DAPI, CD45, HA (using biotinylated Versican G1 HA-binding domain), or SARS-CoV-2; scale bars, 100 μm. **B.** Quantification of SARS-CoV-2 immunostaining in 3 fields of view from matching lung sections. Each point represents an individual mouse. **C.** Quantification of CD45 immunostaining normalized to DAPI stain in 3 fields of view from matching lung sections. Each point represents an individual mouse. **D.** Quantification of HA staining in 3 fields of view from matching lung sections. Each point represents an individual mouse. P values calculated via one way ANOVA followed by Tukey’s HSD test. **E. and F.** Quantification of colocalization between HA (red) and CD45 (green) in 3 fields of view from matching lung sections. Each point represents an individual mouse. **E.** Pearson R and **F.** Mander’s M2 coefficients calculated. P-values calculated with Welch’s two-sample T test and relationships are not significant unless noted otherwise. * = p<0.05

To visualize neutrophil localization in the infected lung, we performed IF with an antibody that binds to the neutrophil marker Ly-6G (Fig 5A). Though there are macrophages that express Ly-6G in the lung after injury [54], they are a very small minority of lung Ly-6G^+^ cells in our MA10 model (S2 Fig). Anti-CD44 treatment reduced the amount of Ly-6G observed (Fig 5B), correlating with the reduction in neutrophils shown via flow cytometry in Fig 2B. Interestingly, colocalization analysis between Ly-6G and HA (Fig 5C-D) showed very similar results to the colocalization between CD45 and HA (Fig 4F-G). This indicates that neutrophils are responsible for at least a significant portion of the observed reduction in CD45 following treatment and are binding via CD44 to the pathological HA matrix in severe COVID-19. Finally, the role of neutrophils in mediating the harmful immune response to MA-10 SARS-CoV-2 was confirmed via depletion of neutrophils through an anti-Ly-6G antibody. Neutrophil infiltration is a hallmark of severe COVID-19, and neutrophil mediation of lung damage and ARDS is well established in pulmonary disease [55]. In this model, the depletion of neutrophils improved survival (Fig 5E), replicating similar experiments with mouse-adapted SARS-CoV-2 in the literature [56,57].

**Fig 5.**
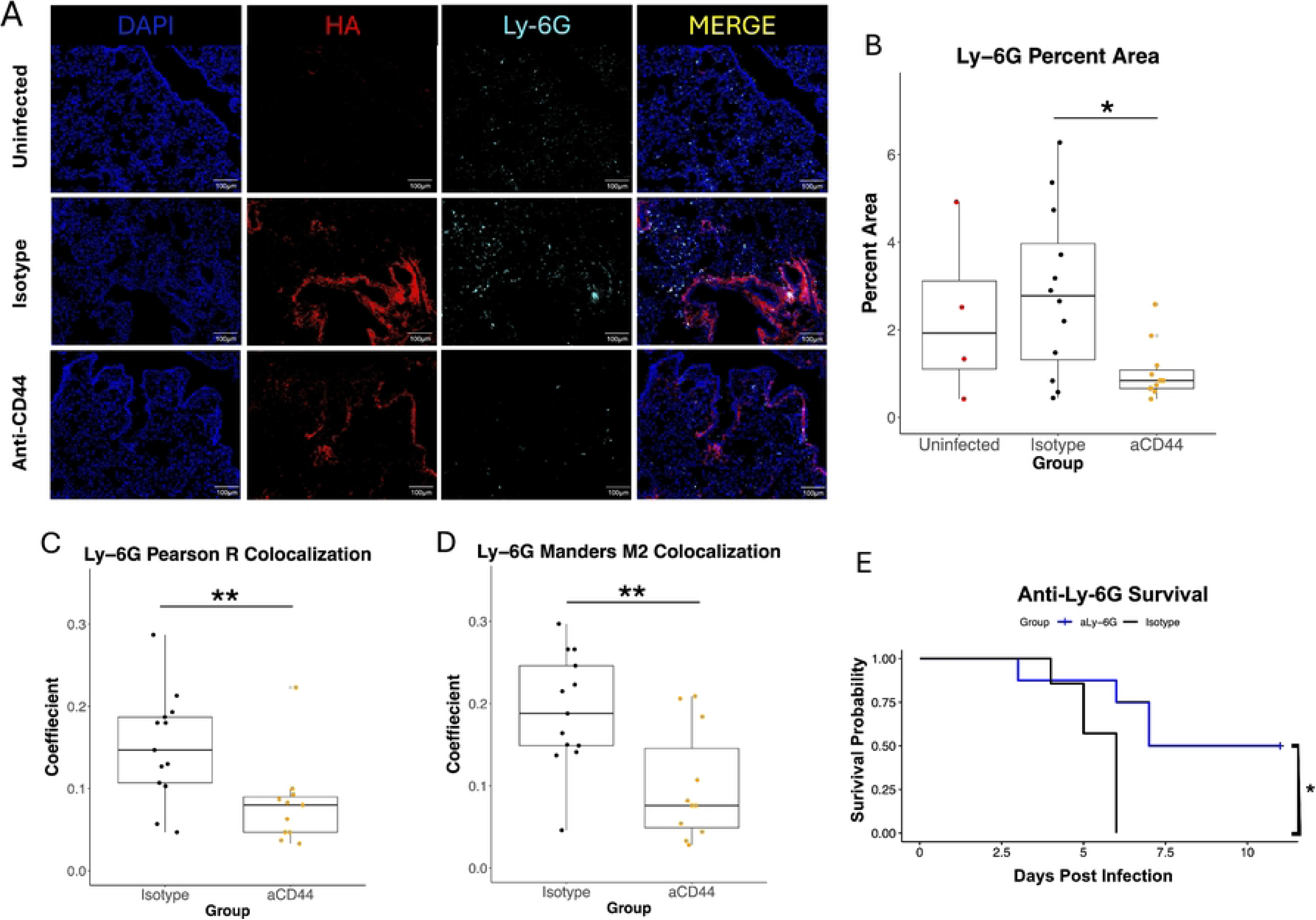
CD44 neutralization decreases colocalization between Ly-6G positive cells and HA. **A.** Immunostaining for DAPI, HA (using biotinylated Versican G1 HA-binding domain), or Ly-6G from superior lung lobe section from 20-week-old MA10-infected C57BL/6 mice treated with 50 µg of two clones of anti-CD44 mAb (aCD44) or an isotype control (Isotype) at day 4 p.i. are presented as representative images. Scale bars, 100 μm. **B.** Quantification of Ly-6G immunostaining in 3 fields of view from matching lung sections. Each point represents an individual mouse. Values calculated via one way ANOVA followed by Tukey’s HSD test. **C. and D.** Quantification of colocalization between HA (red) and CD45 (green) in 3 fields of view from matching lung sections. Each point represents an individual mouse. **C.** Pearson R and **D.** Mander’s M2 coefficients calculated. P-values calculated with Welch’s two-sample T test. N = 12 aCD44-treated mice, N = 13 isotype-treated mice and N = 5 uninfected mice. **E.** Neutrophil depletion via anti-Ly-6G antibody was carried out on 26-week old mice infected with MA-10 SARS-CoV-2. Kaplan-Meier survival analysis was performed. Relationships are not significant unless noted otherwise. * = p<0.05, ** = p<0.01

## Discussion

Here we show that the primary HA receptor CD44 contributes to COVID-19 pathology in a mouse model by mediating neutrophil infiltration/retention into the lungs. Overall, this work provides evidence for a mechanism whereby CD44 on immune cells interacts with the pathogenic HA matrix formed in the lungs during severe COVID-19 to facilitate infiltration of neutrophils, and those cells go on to contribute to a harmful inflammatory response and exacerbate pathology. This represents an advance in our understanding of the role that the extracellular matrix can play during the host immune response to infection.

Our previous work had shown that HA deposition can occur downstream of IL-13 signaling and that a CD44 blockade has positive effects in WT SARS-CoV-2 infection of mice that express human ACE2 [4]. However, we had not explored the effects of a CD44 blockade during infection with a mouse-adapted strain of SARS-CoV-2, which does not cause encephalitis or the same extreme severe disease seen in the K18-hACE2 model [44].

The reduction in lung damage seen in our model (31E-F) is likely the result of the decrease in neutrophil retention after infection and a subsequent decrease in harmful degranulation and release of proinflammatory cytokines. The production of HC•HA likely leads to a pro-adhesive matrix that retains neutrophils in the lungs leading to extensive tissue damage. It is also possible that anti-CD44 treatment blocks neutrophil migration into the lungs during infection. Biochemically cross-linked HA also supports rolling of CD44^+^ cells under much higher shear forces than HA alone, indicating that HC•HA production in blood vessels may increase neutrophil migration [58]. Neutrophil numbers were affected by anti-CD44 treatment, but other cell types such as inflammatory monocytes and T cells were not significantly different between anti-CD44 and Isotype-treated groups despite these other immune cell types increasing with infection and expressing CD44 (Fig 2). This may be due to these cells relying on a CD44-independent mechanism to infiltrate the lung, but there is evidence that neutrophils require HC•HA to bind to fibroblasts, while T cells do not [26].

Our cytokine analysis revealed that many neutrophil chemoattractants were secreted at lower levels in infected mice treated with anti-CD44 (Fig 3A-B). In our previous work we implicated type 2 immune pathways and the cytokine IL-13 as key drivers of COVID-19 pathogenesis upstream of HA deposition [4]. Levels of secreted proteins involved in type 2 immune pathways such as eotaxin (p = 0.017) and IL-5 (p = 0.07) were reduced with anti-CD44 (Fig 3A). These data complement our earlier findings that the induction of HA deposition is downstream of the IL-13-induced type 2 immune response [4].

Alveolar HA is associated with inflammation and severe COVID-19 in humans.[3,5] A reduction in alveolar HA deposition (Fig 4D) provides further evidence that anti-CD44 treatment reduces inflammation in our model. Though the numbers of immune cells are largely unchanged by anti-CD44 treatment as measured by IF (Fig 4C), colocalization of these cells with the HA matrices deposited in the lungs decreased with anti-CD44 treatment (Fig 4E-F). This indicates that anti-CD44 treatment blocks the ability of leukocytes to bind to the cross-linked HA matrix that forms in the infected lung. This provides evidence for a mechanism by which after anti-CD44 treatment not only are these leukocytes not able to traffic efficiently into the inflamed lung effectively, but that if they do enter the lung they cannot bind to the abundant HA (and be retained). Instead, they are free to be cleared before releasing proinflammatory cytokines and contributing to the excess lung inflammation that is a hallmark of severe COVID-19. In Fig 5C-D, we showed that much of this difference in colocalization is due to neutrophils, which was expected based on the flow cytometry data in Fig 2.

While CD44 supports immune cell trafficking during lung disease, it also facilitates the uptake and subsequent degradation of HA into fragments by macrophages [59]. In this way CD44 may contribute to the degradation of HA into proinflammatory LMW fragments that have been seen in the lungs of severe COVID-19 patients, which occurs though an increase in the secretion of HA-degrading enzymes [2]. CD44 has also been implicated as part of the mechanism by which these fragments cause lung epithelial barrier dysfunction. This may be another pathway where blocking CD44 reduces lung damage and the secretion of inflammatory cytokines in our model.

CD44 is the primary HA receptor, but other HA receptors exist and can have roles in the immune response to infection. LYVE-1 and Layilin have been implicated as mediators of immune cell trafficking [60,61]. LYVE-1 is closely related to CD44 [62,63], and has been implicated as the primary HA receptor in endothelia of afferent lymphatic vessels and lymph node sinuses [61]. Layilin is not known to have immune trafficking activity in the lungs and is not structurally related to CD44 or LYVE-1, but it has been shown to mediate lung epithelial barrier function through recognition of HA fragments [64]. These other HA receptors may still contribute to adhesion of leukocytes to the pathogenic HA matrix and their recognition of LMW HA when CD44 is blocked in our model.

Overall, these data provide evidence for a mechanism by which during SARS-CoV-2 a cross-linked HA matrix, composed of HC•HA complexes, binds neutrophils through HA-CD44 interactions, and those neutrophils exacerbate pathology by increasing inflammation and causing lung damage. The finding in this paper that blocking the binding of CD44 to HA improves pathology in a likely neutrophil-dependent manner is relevant to pulmonary immune diseases that are characterized by dysregulated HA matrices such as influenza and RSV.[15,65] Further experiments might include characterizing COVID-19 severity and neutrophil adhesion in our model after blocking HC•HA formation.

## Materials and methods

### Ethics statement

All animal experiments conducted in this study were carried out in accordance with the Animal Welfare Act and the recommendations in the Guide for the Care and Use of Laboratory Animals of the National Institutes of Health. All procedures were approved by the Institutional Animal Care and Use Committee of the University of Virginia (Protocol Number: #4445).

### Virus and cell lines

MA-10 SARS-CoV-2 (BEI: NR-55329) was obtained from the Biodefense and Emerging Infections Research Resources Repository, National Institute of Allergy and Infectious Diseases (NIAID), National Institutes of Health (NIH) [43,66]. The virus was propagated as previously described [4]. The RNA genomes of SARS-CoV-2 MA10 stocks from the second passage were purified with an EZ1 DSP Virus Kit (Qiagen, 62724). Library preparation was performed with a cDNA-PCR Sequencing V14 kit (Oxford Nanopore Technologies, SQK-PCS114) and the cDNA output was sequenced with an Oxford Nanopore MinION. Consensus genome sequences were analyzed using Geneious Prime software (Dotmatics, v.2024.0.5) and demonstrated to possess >99% pairwise sequence alignment with the reference genome for SARS-CoV-2 MA10 (NCBI GenBank accession number MT952602.1).

### Challenge and anti-CD44 treatment

20–24-week-old C57BL/6 mice (Jackson Laboratories) were challenged with 5 x 10^4^ PFU of MA-10 SARS-CoV-2 while under sedation (50mg/kg ketamine; 5mg/kg xylazine). Mice were followed daily for clinical symptoms, which included weight loss (scores 0–5), activity (scores 0– 3), and fur appearance and posture (scores 0–2). Sick mice were euthanized at a humane endpoint of ≥25% weight loss. Mice were given 50µg of each of two anti-CD44 monoclonal antibodies (IM7, BD Biosciences, catalog no. 553131; KM201, SouthernBiotech, catalog no. 1500-01) or 50µg of each of two isotype-matched control antibodies (IgG2b, BD Biosciences, catalog no. 553986; IgG1, Novus Biologicals, catalog no. NBP1-43319-0.5mg) intraperitoneally (i.p.) daily on days 0-3 post-infection.

### Depletion of neutrophils with anti-Ly-6G

26-week-old C57BL/6 mice (Jackson Laboratories) were challenged with SARS-CoV-2 MA10 as described above. Neutrophil depletion was carried out in a manner described previously [56,67]. 100µg of anti-rat Kappa immunoglobulin (MAR18.5, Bioxcell, catalog no. BE0122) was administered i.p. for two days before infection. 50µg of anti-Ly-6G (1A8, Bioxcell, catalog no. BE0075-1) or isotype control (2A3, Bioxcell, catalog no. BE0089) was administered in addition every other day starting the day before infection.

### Western blotting

Right inferior lung samples from day 4 post infection were homogenized in 500µL of PBS, split into two parts, and treated at 37°C for 2 hours; one group treated with 1U Streptomyces hyaluronidase (EMD Millipore, catalog 389561-100U) and one with an equivalent volume of purified water. Laemmli Sample Buffer (Bio-Rad) and 50mM DTT were then added to the samples. Samples were then incubated at 95°C for 5 minutes. Untreated and hyaluronidase treated samples were then loaded to 4-12% Bis-Tris gels and electrophoresis performed at 150 V for 90 mins in MES-SDS running buffer. Proteins were transferred to nitrocellulose membranes at 20 V for 90 mins in 1x transfer buffer (Invitrogen, containing 10% [v/v] methanol). Membranes were then blocked for 1 hour at room temperature (RT) with 10% [w/v] milk-1x phosphate-buffered saline-0.2 % Tween-20 (PBS-T) before incubation with primary antibodies (1:5000, rabbit anti-human polyclonal antibody against IαI [Dako], diluted in 5% [w/v] milk-PBS-T-20) overnight at 4°C on a rocking platform. Membranes were washed in PBS-T before incubation with secondary antibody (donkey anti-rabbit IgG 800Cw; Licor; 1:10,000, diluted in 5% [w/v] milk-PBS-T-20) for 1 h at RT. Blots were imaged on a Licor Odyssey Clx imager (Licor Biosciences GmH).

### Histology

The right superior lobe of the lung was removed on day 4 post-infection. These samples were fixed in formaldehyde and processed before being embedded in paraffin. 5-micron slides were cut and stained with H&E (Thermo Fisher Scientific) using standard protocols. Slides were scanned at 10X magnification. Scoring (1-5) was performed by a blinded independent pathologist using criteria described previously [68].

### Immunofluorescence

Tissues on day 4 post-infection were fixed in 4% PFA for 24 hours at 4°C, then submerged in 30% [w/v] sucrose for one week at 4°C. The tissues were then embedded into frozen blocks, and 5-micron slides were cut. Nonspecific protein was blocked for 1 hour (2% [v/v] donkey serum, 1% [w/v] BSA, 0.05% [v/v] Tween-20) prior to blocking endogenous streptavidin and biotin (Vector Laboratories, catalog SP-2002) for 15 minutes each. Lung sections were then incubated with primary antibodies or Versican G1 HA-binding domain [69] (S1 Fig) overnight, washed in PBS containing 0.05% [v/v] Tween-20, and incubated with secondary antibodies (S1 Fig) for 2 hours at RT followed by 10 minutes of incubation with DAPI (1:1000 concentration) and mounting. Images were captured at 10X magnification with an EVOS FL imaging system (Thermo Fisher Scientific). Blinded analysis of images was performed using Fiji (Version 2.16.0) [70]. The analysis focused on specific regions of interest (airways, vessels, or parenchyma) and a visually determined threshold was applied to each image to capture positive staining while excluding areas with high autofluorescence. Colocalization was done in Fiji using the Coloc2 tool.

### Collection of BALF and flow cytometry

Retro-orbital APC anti-CD45 antibody (1µg in 100µL; Cytek, catalog 20-0451-U100) was administered 5 minutes before mice were euthanized with anesthetic on day 4 post-infection. BAL fluid was collected through cannulation of the exposed trachea. 0.5mL of PBS was flushed first, followed by two flushes with 1mL PBS collected separately. Cells were pelleted, and the supernatant from the first flush was frozen for cytokine analysis. Samples were then stained for 15 minutes at RT with Zombie NIR Fixable Live/Dead (BioLegend, catalog 423106). Cells were then stained with flow cytometry antibodies (S2 Table) for 15 minutes at RT .Samples were then incubated for 30 minutes at 4 °C in BD Cytofix (Cytoperm Fixation/Permeabilization Solution Kit, #555028) for fixation and then were washed and resuspended in FACS buffer. Flow cytometry was performed on a Cytek Aurora flow cytometer (Cytek Bio) and all data analysis performed via OMIQ (OMIQ.ai). The left lobe of the lung was removed and washed in HBSS before being manually diced using scissors and further digested in RPMI 1640 (Gibco) containing 0.17 mg/mL Liberase TL (Roche) and 30 μg/mL DNase (Sigma) for 30 minutes at 37°C. A GentleMACS Dissociator (Miltenyi Biotec) was then used to complete the digestions. Single-cell suspensions were generated by passing samples through a 70µm strainer. Cells were resuspended in FACS buffer before being prepared for flow cytometry in the manner described above.

### Cytokine analysis and ELISA

Cytokine analyses were performed on BALF harvested 4 days post-infection via Luminex Mouse 32-plex (MCYTMAG-70K-PX32, MilliporeSigma). Samples were run following manufacturers protocol after an 18-hour incubation before being run on Luminex analyzer (MAGPIX). Albumin concentration was measured on BALF diluted 1:1000 from day 4 post-infection using an ELISA kit (Abcam). Data was pre-processed and visualized using R software. Cytokine values were log-transformed, centered, and scaled before being visualized as a heatmap using the pheatmap package in R.

## Acknowledgements

The authors thank the UVA Flow Cytometry Core Facility, the UVA Research Histology Core, and the UVA Biorepository and Tissue Research Facility. The authors thank Mary Young and Dr. Mehmet Tanyüksel for their support, and Amy Mathers, Katie Barry, and Chava Castaneda Barba for guidance on sequencing SARS-CoV-2 MA10 stocks. Fig 1A was created in BioRender (Hart, D. (2025) https://BioRender.com/591ldies).

This work was supported by the UVA Infectious Disease Training Grant (5T32AI007046-48), the Manning Family Foundation, the US National Institutes of Health (R01 1R01HL171283-01A1), the Wellcome Trust (Discovery Award 304200/Z/23/Z ), and Robert and Dr. Lisa Henske.

## Supplemental Material

**Supplementary Figure S1. Anti-CD44 monoclonal antibody treatment does not reduce IV^+^ neutrophil numbers.** Changes in whole lung IV^+^ lymphocyte numbers in whole lung homogenates taken from MA10-infected mice on day 4 post infection. Neutrophils (CD45^+^ CD11b^+^ CD11c^-^ Ly6C^hi^ Ly6G^+^), Inflammatory monocytes (CD45^+^ CD11b^+^ CD11c^-^ Ly6C^hi^ Ly6G^-^), B cells (CD45^+^ CD19^+^ SSC^lo^), CD8 T cells (CD45^+^ CD3^+^ CD8^+^ SSC^lo^), and CD4 T cells (CD45^+^ CD3^+^ CD4^+^ SSC^lo^) are shown. No significant differences in P values calculated via one way ANOVA followed by Tukey’s HSD test.

**Supplementary Figure S2. Neutrophils make up the vast majority of Ly-6G^+^ cells.** Neutrophils (CD45^+^ CD11b^+^ CD11c^-^ Ly6C^hi^ Ly6G^+^) as percentage of all live Ly-6G^+^ cells in whole lung homogenates taken from MA10-infected mice on day 4.

**S1 Table. Immunofluorescence antibodies**

**S2 Table. Flow cytometry antibodies**

